# ^2^H MRI-based quantification of leucine uptake in glioblastoma multiforme

**DOI:** 10.1101/2025.04.23.648848

**Authors:** Samantha McClendon, Xia Ge, Kyu-Ho Song, Dustin Grief, Shannon P. Fortin Ensign, Vikram D. Kodibagkar, Leland S. Hu, Joel R. Garbow, Scott C. Beeman

## Abstract

Glioblastoma (GBM) brain tumors are among the most lethal of all human cancers, with a median survival time of ∼15 months. Treatment planning requires radiologic demarcation of tumor boundaries with contrast-enhanced magnetic resonance imaging (CE MRI); however, significant tumor burden extends beyond the contrast-enhancing margins of the tumor. GBM tumors have an increased expression of amino acid (AA) transporters, including the Alanine, Serine, Cysteine Transporter 2 (ASCT2) and the L-Type Amino Acid Transporter 1 (LAT1). This upregulation has been leveraged in positron emission tomography (PET) studies to detect tumor burden beyond the contrast-enhancing margins identified by standard-of-care CE MRI. Here we leverage recent approaches in deuterium metabolic magnetic resonance with this known upregulation of AA transporters in GBM to demonstrate that ^2^H MR can detect glioma based on enhanced branched-chain amino acid (BCAA) uptake. To the best of our knowledge, these data represent the first non-invasive quantification of AA concentrations in brain tumor and raises the potential to (i) detect tumor burden beyond contrast-enhancing margins and (ii) quantify AA metabolism using ^2^H MR spectroscopy.

## 1. Introduction

Glioblastoma (GBM) brain tumors are among the most aggressive and lethal of all human cancers, with a 5-year survival of less than 5% and median survival time of ∼15 months.^1,2^ Standard-of-care surgery and radiation therapy rely on accurate detection and demarcation of these tumors, most commonly achieved via contrast-enhanced magnetic resonance imaging (CE MRI). CE MRI is the cornerstone of nearly all radiologic and clinical management of GBM and relies on the leakage of a gadolinium-based contrast agent through the permeable blood brain barrier (BBB) that is characteristic of tumor tissue. As this method is based on the extravasation of the agent from the vascular space into the interstitial space, it does not directly identify tumor cells, which often extend well beyond the boundaries of BBB disruption.^3,4^

GBM cells are known to have reprogrammed metabolism of amino acids (AAs), which aids in their proliferation, signal transduction, adaptation, and survival.^5^ Increased uptake of AAs is facilitated by upregulated expression of amino-acid transporters including the Alanine, Serine, Cysteine Transporter 2 (ASCT2) and the L-Type Amino Acid Transporter 1 (LAT1).^6–11^ The known upregulation of AA transporters in GBM has enabled positron emission tomography (PET) studies to detect tumor based on increased uptake of radiolabeled AA tracers. Importantly, because AAs are actively transported directly into GBM cells, these radiotracers can identify tumor cells well beyond the boundaries of BBB disruption.^12–14^ Many variations of AA-PET exist, most commonly using radiotracers O-(2-[18F]-fluoroethyl)-L-tyrosine (FET),^6–8,12–23,24(p11),25^ L-methyl- 11C-methionine (MET),^6–8,12–23,24(p11),25^ and 3,4-dihydroxy-6-[18F]fluoro-L-phenylalanine (FDOPA).^6–8,12–23,24(p11),25^ The branched-chain amino acid (BCAA) leucine-analog 18F- Fluciclovine, which is FDA-approved for imaging/detection of suspected prostate tumors,^26^ has been used to image GBM in rodents and is positively correlated with tissue AA transporter levels.^27^

However, AA-PET has major limitations in accessibility: (i) short half-lives of radio-labelled AA analogues necessitate imaging proximal to one of approximately 250 cyclotron facilities in the country,^28^ typically limiting AA-PET access to major metropolitan regions; (ii) PET introduces clinical challenges in scheduling, time to diagnosis, and medical costs and, importantly, additional burden on patients who need to schedule a PET scan in addition to routine MRI; (iii) AA-PET tracers lack FDA approval for brain tumor imaging and thus remain non-reimbursable and inaccessible for most patients. Thus, most GBM patients never undergo this type of imaging. A method for detecting GBM cells based on AA uptake, but with accessibility like that of CE-MRI, would be a significant advance.

Recent studies show that replacing protium with deuterium in metabolically active molecules such as glucose and acetate allows quantification of their uptake using ^2^H metabolic magnetic resonance (^2^H metabolic MR).^29–34^ 2H metabolic MR, including both ^2^H imaging and ^2^H single-voxel spectroscopy, is a method for measuring the uptake and metabolism of deuterium-enriched substrates. Among the many deuterated substrates incorporated into ^2^H metabolic MR experiments are glucose, acetate, choline, fumarate, methionine, water, and recently, alanine.^35,36^ Deuterated glucose (D-[6,6^’^ - ^2^H_2_] glucose) has been used extensively to study bacterial metabolism,^37^ retinal metabolism,^38,39^ glycogen synthesis,^40^ red blood cell metabolism,^41^ and, notably, cancer metabolism as related to the Warburg effect.^29,42^

Motivated by advances in AA-PET and ^2^H metabolic MR, we test the hypothesis that enhanced uptake of deuterated AAs in GBM can be quantified with ^2^H MR and demonstrate this concept, which we refer to as ^2^H AA-MR, in a rodent model. The success of this proof-of-concept study lays the foundation for a novel, accessible tool to directly image GBM cells in the brain independent of BBB disruption.

## 2. Methods

### In vivo tumor and d_10_-leucine dose preparation

All animal experiments were approved by the Washington University Institutional Animal Care and Use Committee. Three tumor-bearing mice were prepared by stereotactically injecting 50k GL261 cells into the dorsal striatum of 70-day-old female C57BL/6 mice. Tumors were allowed to grow for approximately 20 days, reaching volumes of 58, 49, and 46 mm^3^, respectively. A 100 mM d_10_-leucine solution was prepared by adding DL-leucine-d_10_ (98 atom % D) (Sigma-Aldrich, Co., St. Louis, MO) to deionized water and sonicating in an ultrasonic bath until dissolved. Note that in a separate experiment without an imaging component, ^2^H leucine doses of up to 70.6 mg/100 g, were found to be safe in mice, with four female, 10-month-old, C57BL/6 mice surviving a 1 mL dose of 200 mM leucine (data not shown).

### Time course ^2^H MR Spectroscopy (MRS)

MRS was completed for three tumor-bearing and three control mice using a home-built volume transmit, surface receive coil configuration on a 11.7 T Varian system. For each animal, two pre-injection, natural-abundance ^2^H Spin-Echo full-Intensity Acquired Localized (SPECIAL) spectra (5 minutes each, TR/TE = 450/4.27 ms) ^43^ were collected prior to ^2^H leucine administration. Next, a target dose of 500 µL of 100 mM d_10_-leucine solution was injected *via* a subcutaneous catheter between the shoulder blades of the dorsal surface region (total dose 35.3 mg/100g) while the animal was in the scanner. Immediately following injection, ^2^H SPECIAL spectra were collected in 5-minute increments for 70 mins post injection: TR/TE = 450/4.27 ms. Voxels were chosen to maximize volume within tumoral regions (3.3 × 4.9 × 4 mm^3^, 4.2 × 3.7 × 5 mm^3^, 4 × 4 × 4 mm^3^). For healthy animals, a 3 × 6 × 5 mm^3^ voxel was placed within the brain to maximize SNR.

### Cell culture and extraction

GBM patient-derived xenograft (PDX) cell line GBM22 was provided by the Mayo Clinic Brain Tumor Patient-Derived Xenograft National Resource, Phoenix, AZ, USA.^44^ Cells were first plated in DMEM 10% FBS and then serum starved (DMEM 0.1% BSA) for 12 hours. Next, cells were cultured in either DMEM 5% FBS or DMEM 5% FBS supplemented with 25 mM DL-Leucine-d_10_ (98 atom % D) (Sigma-Aldrich, Co., St. Louis, MO) for 24 hours. Cells were harvested and rinsed twice with PBS to ensure removal of external media prior to extraction. Metabolites were extracted by a modified acetonitrile extraction protocol,^45^ with basic steps as follows: (i) Cell pellets of ∼10 million cells were homogenized in a solution of acetonitrile:isopropanol:water (volume 3:3:2), (ii) Resulting cell extract and debris were centrifuged at 10,000 xg and 4 °C for 30 minutes, (iii) Recovered supernatants were roto-evaporated to dryness, (iv) Dried samples were reconstituted in a solution of acetonitrile: water (volume 1:1), then centrifuged and roto-evaporated again, and (v) Dried samples were reconstituted in 200 µL deionized water (Advanced Biomatrix, Inc. MS) and transferred to 3 mm tubes for NMR.

### ^2^H-NMR spectroscopy

^2^H NMR was performed on a Bruker Advance III NMR console at 9.4 T. The broadband channel of a Bruker 5 mm VSP NMR probe ([^109^Ag-^31^P]/^13^C/^1^H with ^2^H lock and Z gradient) was tuned to ^2^H, at a resonant frequency of 61.42 MHz, for direct detection of the NMR signal. Spectra were collected with a 90° pulse, relaxation delay of 1s, acquisition time of 2s (recycling time of 3s), and sweep width of 3076.92 Hz. 5088 scans were acquired for each experiment.

### Phantom Preparation

A d_10_-leucine phantom of 20 mM was created in a 1.5 mL centrifuge tube to measure relaxation time constants. The phantom was prepared by dissolving DL-Leucine-d_10_ (98 atom % D) (Sigma-Aldrich, Co., St. Louis, MO) in a 2% low gelling temperature agarose solution (Sigma-Aldrich, Co., St. Louis, MO) heated on a hot plate, with D_2_O added to enhance the 4.7 ppm resonance. The sample was transferred to a 1.5 mL centrifuge tube and allowed to cool until gelled.

### T1 and T2 quantification

Longitudinal and transverse relaxation time constants (T1 and T2, respectively) were measured using the d_10_-leucine gel phantom on a Varian 11.7 T small animal MRI system with a custom-built ^2^H coil. T1 values of the HOD resonance and (-CD_3_)_2_, -CD_2_, gamma-D, and alpha-D resonances of d_10_-leucine were measured with a two-pulse inversion-recovery experiment. Forty exponentially-spaced inversion times ranging from 0.02 to 2 s were used, with a TR of 2500 ms. A Carr-Purcell-Meiboom-Gill (CPMG) spin echo train experiment was then used to quantify the T2 values of the same resonances; 40 exponentially-spaced echo times ranging from 0.005 to 0.75 s were used, with a TR of 2500 ms.

### Data analysis

MRS frequencies, amplitudes, and relaxation time constants were estimated using the Bayesian Analysis Toolbox, a data analysis package developed at Washington University in St. Louis.^46,47^ Frequencies and decay rate/time constants across all animals and spectra in the time series were estimated jointly (i.e., all frequencies and decay rate constants are assumed to be the same across animals and points in the MRS time series), and all amplitudes across time points and animals were estimated independently. Deuterated leucine concentrations were calculated by referencing (-CD_3_)_2_ peak amplitudes to natural-abundance HOD (average of two 5-min ^2^H SPECIAL scans, referenced to a concentration value of 16.35 mM ^48^) and adjusting for T1, T2, and stoichiometry/label loss.^49,50^ Deuterated leucine concentrations from cell extraction experiments were calculated likewise, referencing the (-CD_3_)_2_ peak amplitude to a natural abundance HOD concentration of 15.37 mM, as measured and calculated for the deionized water solvent used for NMR (Advanced Biomatrix, Inc. MS) (see NMR experiment in Supplemental Figure 1). For relaxation experiments, T1 and T2 time constants were quantified by fitting a standard three-parameter (mono-exponential plus a constant) model to the data.

### Statistics

For *in vivo* data, a Student’s t-test was used to compare deuterated leucine concentrations for the tumoral and control groups at the 70-minute time point (*α* = 0.05).

## 3. Results

### *In vivo* ^2^H AA-MR

*In vivo* experiments support the feasibility of ^2^H AA-MR for quantifying uptake of AAs in GBM. Enhanced uptake of d_10_-leucine in tumor tissue was observed over a 70- minute time course (Figure 1(a)), with ∼4-fold enhancement (p = 0.001, based on 70-minute time point) compared with comparable tissue in control animals observed at the 70-minute time point and a greater than 4-fold increase over the time course, compared to natural abundance ^2^H leucine (conservatively estimated to be at the level of the noise floor) (Figure 1(b)).Tumoral concentrations of 2.0 mM ± 0.3 mM (at 70 min.) were achieved using doses of 35.5 mg/100 g, while essentially no uptake was seen in normal-appearing brain. Leucine concentrations in healthy brain were largely below the noise floor (around 0.5 mM; note that data points beneath the noise floor reflect the Bayesian probability software’s best estimate but are not deemed quantifiable by the authors). Assuming an approximate linear uptake of d_10_-leucine over the first 45 minutes of data collection, we estimate an uptake rate of approximately 2.8 μg/min. A corresponding plot of HOD concentrations shows no difference between tumor and control (p = 0.77, based on 70-minute time point) and an increase to approximately 1.5 times natural abundance HOD over the 70-minute time course in *both* tumoral and control animals (p < 0.001, Figure 1(c)).

**Figure 1.**
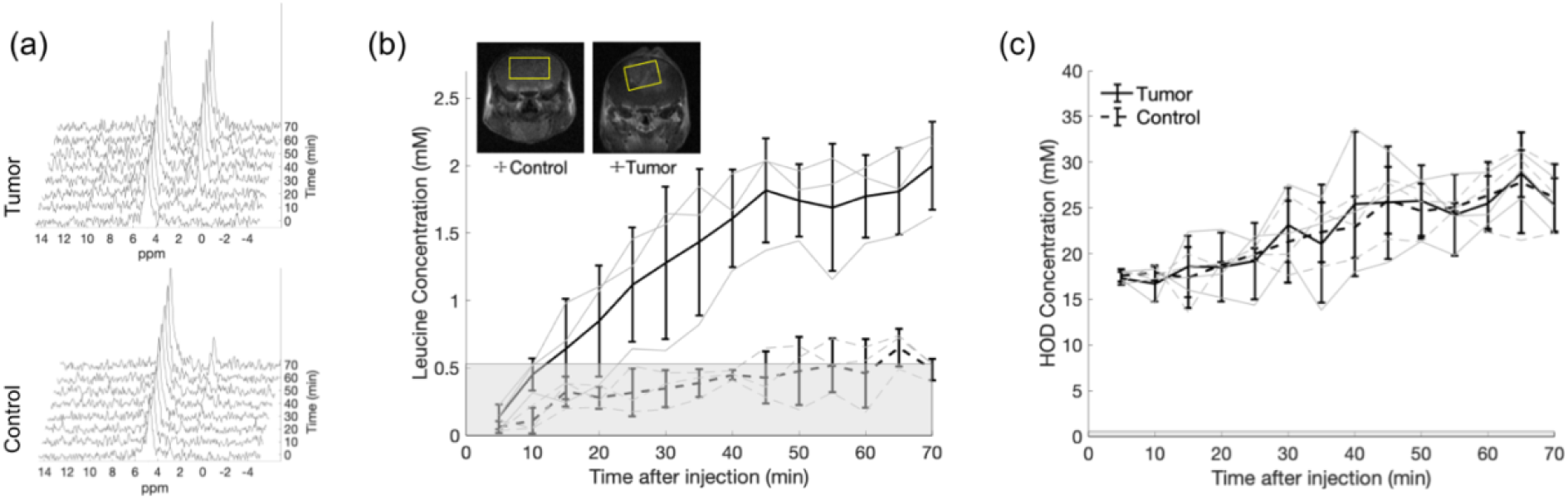
(a) SPECIAL ^2^H MR spectra of voxels representing tumor and control tissue (see voxel placement in (b)) over 70 minutes post-subcutaneous injection (5-minute intervals, lb = 10 Hz) show an increase in the leucine-associated peak at ∼0.7 ppm. (b) Concentrations of ^2^H leucine over the 70-minute time course, calculated from (CD_3_)_2_ peak amplitudes referenced to the natural- abundance HOD peak. (c) Concentrations of HOD over the 70-minute time course, again referenced to the natural-abundance HOD peak. For each plot, light grey lines represent individual animals, while bolded lines represent averages across animals. Concentrations below the noise floor are shaded in grey.

### ^2^H NMR of cell extracts

The spectra generated from d_10_-leucine-exposed cells show ^2^H leucine-associated resonances: (-CD_3_)_2_ at 0.78 ppm, and (-CD_2_) and γ-D at 1.5 ppm. Given the extraction procedure used here (see Methods), we measure a ^2^H leucine concentration of 0.45 mM for these extracts. In contrast, control spectra from cells not exposed to d_10_-leucine but undergoing the same extraction procedure do not exhibit any leucine-associated resonances above the noise floor but instead only exhibit the expected natural-abundance HOD resonance, referenced at 4.7 ppm (Figure 2).

**Figure 2.**
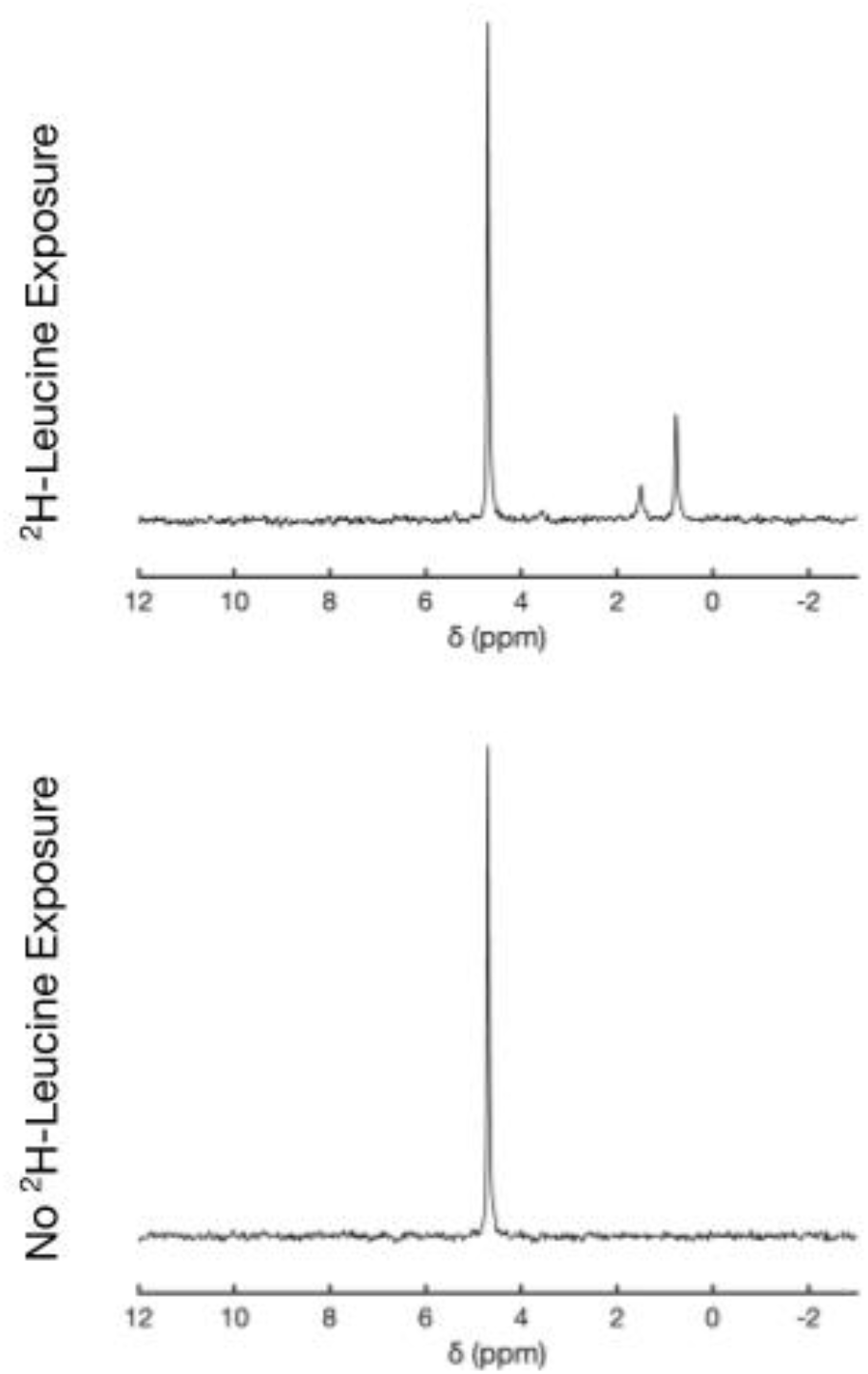
Comparison of ^2^H NMR spectra (5088 scans) of GBM 22 cell extracts from d_10_-leucine- exposed and non-exposed cells, showing leucine-associated (-CD_3_)_2_ resonance (0.78 ppm) and (- CD_2_) and γ-D resonance (1.5 ppm) for ^2^H-leucine-exposed cells. The natural-abundance HOD resonance is referenced to 4.7 ppm.

### *In vitro* relaxation time constant (T1/T2) measurements

Measured T1 and T2 relaxation time constants were used in calculations of d_10_-leucine concentrations. Inversion recovery and CPMG experiments were performed using a 20 mM d_10_-leucine phantom and standard (three-parameter) mono-exponential-plus-a-constant models were fit to the data to determine T1 and T2 time constants. Leucine-associated resonances and their T1 and T2 relaxation time constants at a concentration of 20 mM were: (-CD_3_)_2_ T1/T2 = 166/148 ms, (-CD_2_) and γ-D 117/60 ms, α-D 134 /46 ms. For the HOD peak, T1/T2 = 393/40 ms (Figure 3).

**Figure 3.**
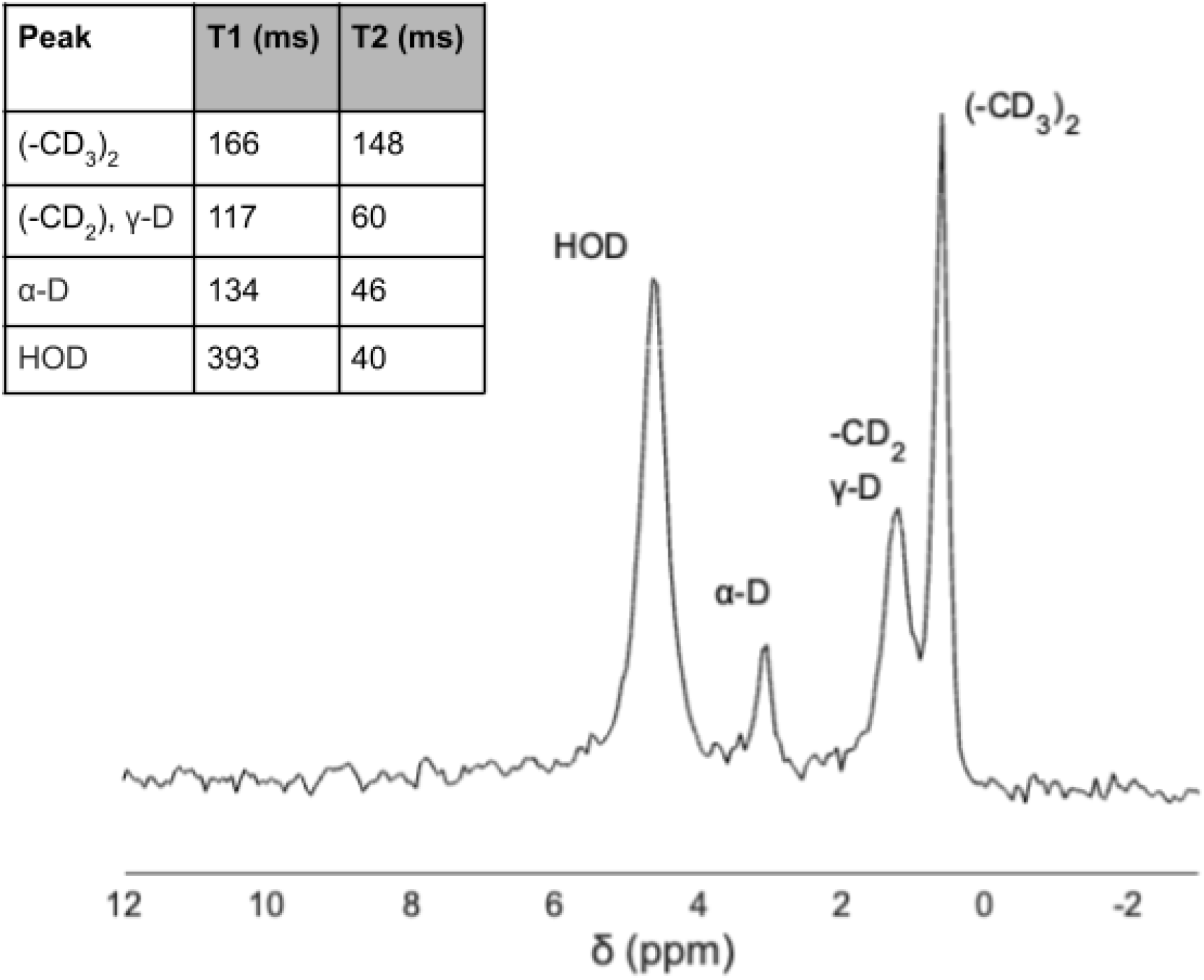
SPECIAL ^2^H MR spectrum (4 × 4 × 4 mm^3^ voxel) of a ∼20 mM deuterated leucine phantom in 2% agarose obtained at 11.7 T, plotted with 10 Hz line broadening. (The HOD resonance was enhanced with addition of D_2_O to the phantom.) Measured T1 and T2 values are tabulated in the insert in the upper left of the figure.

## 4. Discussion

This study provides proof-of-concept data to support a ^2^H AA-MR method to quantify the preferential uptake of AAs in GBM vs normal brain. We show a ∼6-fold enhancement in ^2^H leucine concentrations in tumoral compared to healthy tissue following a subcutaneous injection of the agent; tumor concentrations of upwards of ∼2 mM were achieved using doses of 35.5 mg/100 g. Notably, ^2^H NMR of cell extract experiments, in which we exposed cultured GBM cells to ^2^H leucine and used ^2^H NMR to quantify intracellular leucine concentration after washing cells and extracting metabolites, provide direct evidence of cellular leucine uptake independent of vascular leakage. This idea is further supported by the similarity in HOD time-series profiles between tumoral and healthy tissue (Figure 1(c)). HOD is produced via systemic metabolism of injected ^2^H leucine and, being freely diffusible, enters the blood stream and distributes through the body in a way that can be approximated as a slow infusion of HOD (a similar delivery mechanism to the subcutaneous delivery of ^2^H leucine). The similarity in the HOD time-series profiles between tumoral and healthy tissue suggests that delivery mechanisms that are not cell membrane transporter mediated are similar between tumor and healthy tissue. Given our in vitro data and these control HOD time-series data, we thus conclude that the difference between tumoral and healthy brain ^2^H leucine concentrations is likely mediated by transporter-mediated cellular uptake of ^2^H leucine into tumor cells. This idea of direct AA uptake is also well supported by extensive literature supporting the transporter-facilitated affinity of GBM to circulating leucine.^6–11^ While further exploration is necessary to: (i) understand the *extent* of uptake versus possible extravasation of ^2^H leucine and (ii) resolve the effects of differing cell types on degree of uptake, the leucine uptake confirmed by these ^2^H NMR experiments confirms the potential for direct ^2^H leucine uptake in GBM *in vivo*.

Beyond ^2^H AA uptake, the topic of this study, the ^2^H AA-MR method may provide an opportunity to explore AA metabolic activity in brain tumors. This future direction could be a significant development in tumor diagnosis and staging, as abnormal AA metabolism is a critical biomarker of GBM malignancy. For example, enhanced GBM uptake and catabolism of BCAAs to lactate have been shown to drive GBM cell proliferation and are associated with a particularly aggressive hypoxic GBM phenotype.^10,11,51^ By mimicking ^2^H metabolic MR studies that track the appearance of glutamine/glutamate and lactate peaks in ^2^H NMR spectra following administration of deuterated glucose, we might quantify the downstream appearance of known leucine metabolites such as glutamate/glutamine, lactate, or acetoacetate. However, doing so *in vivo* would require the ability to detect signal from peaks associated with lower ^2^H concentrations than that of the (CD_3_)_2_ peak of d_10_-leucine here, as downstream metabolites would: (i) have fewer potentially deuterated functional groups, (ii) have probabilistically fewer deuterons in those locations, and (iii) potentially be more transitory in nature. In this regard, it must be noted that dramatic improvements in SNR have been made in the short period since the introduction of ^2^H metabolic MR that have not been leveraged in this study. For example, Peters, et al., have demonstrated a ^2^H multi-echo balance steady-state free-precession (ME-bSSFP) that improved SNR by a factor of ∼2-5 (depending on condition and spectrum peak).^52^ Expanding on this work, Montrazi, et al., recently demonstrated a weighted average ^2^H chemical shift imaging steady-state free precession (CSI- SSFP) sequence that increases SNR by a factor of ∼5 over the already-improved ^2^H ME-bSSFP, for a total 10-25x improvement over the original ^2^H CSI sequence used by De Feyter, et al ^29,53^ and the ^2^H NMR experiments used herein. Additionally, in a separate study not reported here, the use of higher ^2^H leucine doses of up to 70.6 mg/100 g were found to be safe in mice (see Methods), potentially doubling the ^2^H leucine signal available in future studies. Combined, these methods could provide a theoretical 20-50 fold increase in detectability, and even if the gains are more modest, significant improvement in signal is expected.

In this proof-of-principle study, we report an uptake rate assuming linear uptake, a rough but reasonable approximation based on the data. This lays a foundation for future studies focused on appropriate kinetic modelling, accounting for blood concentration and potential downstream metabolic conversion, as demonstrated for example in D-[6,6^’^ - ^2^H_2_] glucose ^2^H metabolic MR.^54^ While subcutaneous injections were used here as a surrogate for constant ^2^H leucine administration to the bloodstream, preliminary data using IP and IV administration of the deuterated leucine produce comparable results (Supplemental Figure 2), suggesting consistency across dosing schemes. IV administration may allow for more control of dose timing and blood substrate levels in dynamic uptake and clearance studies.^55^

A noninvasive and accessible method for quantifying AA uptake would be a significant clinical advance. Primarily, ^2^H AA-MR provides an opportunity for imaging GBM directly based on enhanced expression of AA transporters, rather than BBB disruption. Practically, this method may readily translate to the clinic. Although x-nuclei capabilities are at present more commonly used in academic hospital settings, modification of clinical MRI systems to provide x-nuclei capabilities can be accomplished on-site with appropriate radiofrequency coils, filters, and radiofrequency amplifiers and pre-amplifiers.^56^ PET is currently more commonly available in the clinic; however, these ^2^H MRI modifications are highly affordable compared to the installation of PET imaging and cyclotron facilities. Contemporary coil design also allows for simultaneous ^1^H and ^2^H scanning, further reducing scan time per patient. Simultaneous ^1^H/^2^H scanning would fit well into a clinical workflow at many sites while eliminating the significant scheduling, cost, and quality-of-life burden of PET imaging.

## 5. Conclusions

Herein, we present the first proof-of-principle confirming that ^2^H MR can be used to quantify enhanced AA uptake by GBM. This ^2^H MR method has the potential to significantly advance accurate and accessible imaging of these aggressive, infiltrative brain tumors based directly on AA uptake into GBM cells, rather than BBB breakdown. Moreover, the preliminary success of this method provides a foundation for the study of differential AA metabolism in GBM.

## List of Abbreviations

^2^H AA-MR: Deuterium amino acid metabolic resonance
AA: Amino acid
ASCT2: Alanine, Serine, Cysteine Transporter 2
BBB: Blood brain barrier
BCAA: Branched-chain amino acid
CE MRI: Contrast enhanced magnetic resonance imaging
FDOPA: 3,4-dihydroxy-6-[18F]fluoro-L-phenylalanine
FET: O-(2-[18F]-fluoroethyl)-L-tyrosine
GBM: Glioblastoma multiforme
LAT1: L-Type Amino Acid Transporter 1
MET: L-methyl-11C-methionine
PET: Positron emission tomography
SPECIAL: ^2^H Spin-Echo full-Intensity Acquired Localized

## Acknowledgements

Grant acknowledgements: The research presented was partly supported by National Institutes of Health grant R01-EB032376, Arizona Biomedical Research Commission grant RFGA2022-010, and the ASU-Mayo Alliance Seed Grant.

Special thanks to Joseph Ackerman (Washington University, St. Louis, MO) for his insights during experimental planning and continued valuable feedback throughout the course of this project.

## Data Availability

The data that support the findings of this study are available from the corresponding author upon reasonable request.

## Notes

### Competing Interest Statement

Author Leland Hu is a paid speaker and serves on the advisory board for Bayer, a company which produces oncology pharmaceuticals.
No other authors have conflicts of interest to disclose.

