## Supplemental Figures for "^2^H MRI-based quantification of leucine uptake in glioblastoma multiforme"

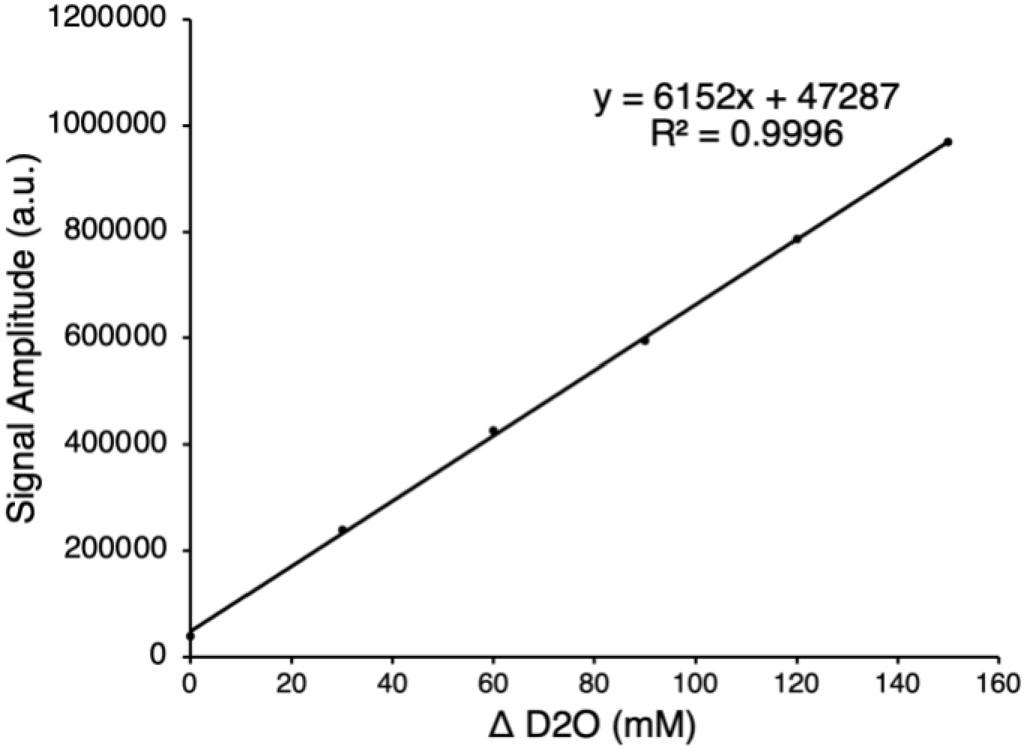


**Supplemental Figure 1:** Signal amplitudes (arbitrary units) from $^2$H NMR spectra plotted for a series of samples of deionized water (Advanced Biomatrix, Inc. MS) with increasing $^2$H concentration achieved by adding $D_2O$. Natural abundance $^2$H was determined as 15.37 mM.

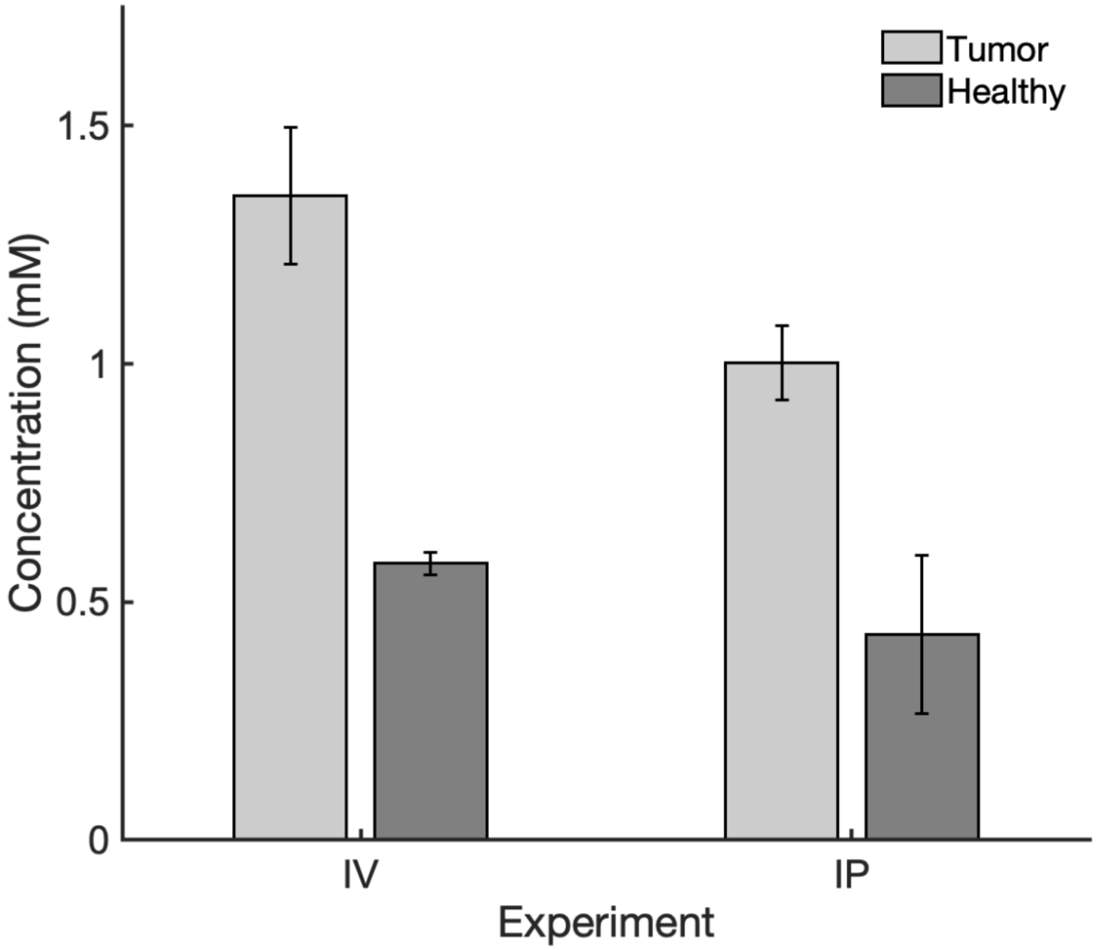


**Supplemental Figure 2:** SPECIAL $^{2}$H MR-measured *in vivo* concentrations of $d_{10}$-leucine following IV or IP leucine administration (1 animal each; Error bars represent standard deviation across three consecutive spectra). Dosing schemes were as follows: Three IV bolus injections of 300 µL of 100 mM $d_{10}$-leucine in saline (pH, ~7; total dose = 12 mg/100 g) or one IP bolus injection of 900 µL of this same solution. Animals were allowed to recover for one hour prior to MR spectroscopy. Three serial $^{2}$H SPECIAL spectra (15 min of data acquisition each) were collected *in vivo*: TR/TE = 250/4.27ms; 3600 averages each.
